# Native and engineered clifednamide biosynthesis in multiple *Streptomyces* spp

**DOI:** 10.1101/197616

**Authors:** YuncI Qi, Edward Ding, Joshua A.V. Blodgett

## Abstract

Polycyclic tetramate macrolactam (PTM) natural products are produced by actinomycetes and other bacteria. PTMs are often bioactive, and the simplicity of their biosynthetic clusters make them attractive for bioengineering. Clifednamide-type PTMs from *Streptomyces* sp. JV178 contain a distinctive ketone group, suggesting the existence of a novel PTM oxidizing enzyme. Here, we report the new cytochrome P450 enzyme (CftA) is required for clifednamide production. Genome mining was used to identify several new clifednamide producers, some having improved clifednamide yields. Using a parallel synthetic biology approach, CftA isozymes were used to engineer the ikarugamycin pathway of *Streptomyces* sp. NRRL F-2890 to yield clifednamides. Further, we observed that strong CftA expression leads to the production of a new PTM, clifednamide C. We demonstrate the utility of both genome mining and synthetic biology to rapidly increase clifednamide production and identify a PTM tailoring enzyme for rational molecule design.

---

Actinomycete bacteria are widely studied for their ability to produce diverse bioactive secondary metabolites. Already the source of nearly two-thirds of clinical antibiotics^1^, actinomycete genome sequencing has revealed a wealth of previously unrecognized biosynthetic gene clusters^2–4^. With many of these clusters apparently encoding drug-like molecules, these organisms remain as promising sources of much-needed future antibiotics and other therapeutics^5,6^.

An unusually high proportion of sequenced actinomycete genomes contain polycyclic tetramate macrolactam (PTM) biosynthetic clusters^7^. PTMs are of therapeutic interest, with family members having documented activity against bacteria, protozoa, fungi, plants, and cancer cell lines^8–11^. In addition to their bioactivity, the relative simplicity and commonality of PTM biosynthetic loci has made them attractive targets for genomics-based discovery and engineering via synthetic biology approaches^12–14^. Despite containing only 3-6 genes, these small clusters encode diverse structures^15^ (Figure 1).

**Figure 1.**
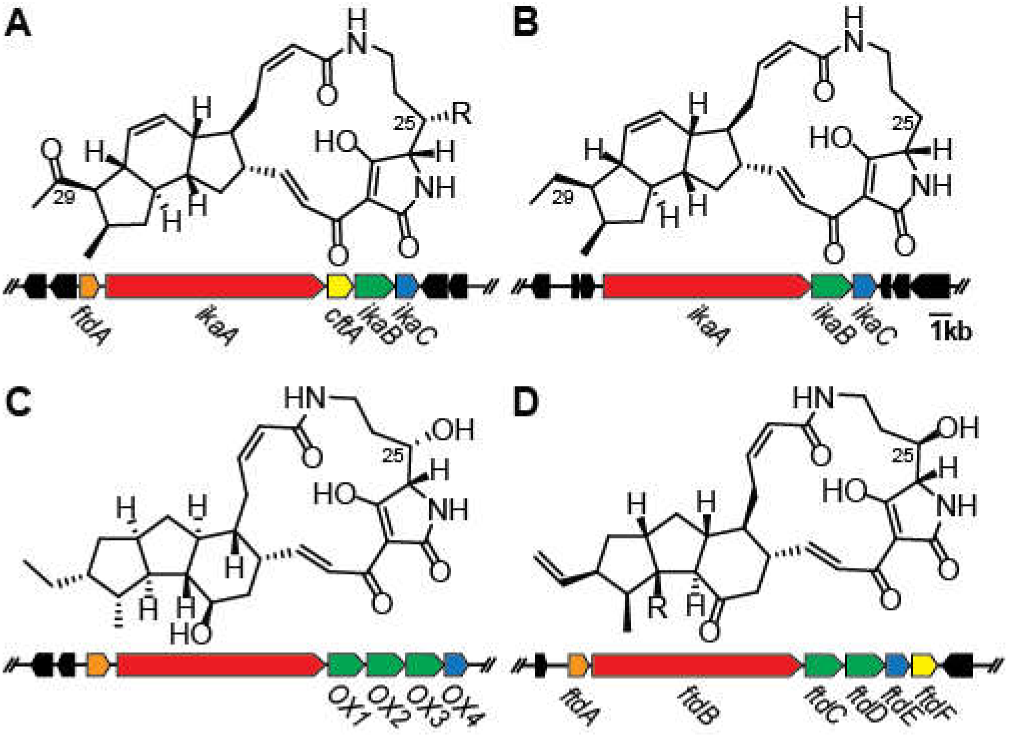
Representative PTM biosynthetic clusters and their products. (A) the *Streptomyces* sp. JV178 clifednamide cluster (A: R=H, B: R=OH); (B) the *Streptomyces* sp. ZJ306 ikarugamycin cluster; (C) The *Lysobacter enzymogenes* C3 HSAF cluster; (D) the *Streptomyces* sp. SPB78 frontalamide cluster (A: R=OH, B: R=H). Orange ORFs encode for sterol desaturases, red for iterative polyketide synthase non-ribosomal peptide synthase fusion proteins, green for FAD-dependent oxidoreductases, blue for zinc-dependent alcohol dehydrogenases, and yellow for cytochrome P450s. ORFs in black are not conserved between PTM biosynthetic clusters.

The clifednamides are a family of PTMs discovered from *Streptomyces* sp. JV178, an environmental isolate from Connecticut garden soil^16^. The clifednamides have therapeutic potential due to their structural similarity with ikarugamycin, which is active against a wide range of organisms^8,10,11^. The clifednamides are distinguished from ikarugamycin by a ketone group on carbon 29 (Figure 1A). Despite growing research interest in PTMs, no structure-function studies have been carried out thus far. Comparing the bioactivities of the long-studied ikarugamycin and its clifednamide analogs could provide valuable insight. However, efforts to extensively profile clifednamide bioactivities have been limited by low yields from JV178 (Table 1).

**Table 1.** PTM titers from key strains in this study

| Strain | Ikarugamycin | Clifednamide A | Clifednamide B | Clifednamide C |
| --- | --- | --- | --- | --- |
| *S. sp.* JV178 | Undetected | 0.94 μM | 1.87 μM | Undetected |
| *S. sp.* NRRL F-2890 | 28.2 ± 5.27 μM | Undetected | Undetected | Undetected |
| *S. sp.* NRRL F-2890<br>*PermE* * <sub>pDA1652</sub> - *cftA* <sub>JV178</sub> | 1.10 ± 0.68 μM | 13.73 ± 1.04 μM | Undetected | 2.62 ± 0.70 μM |
| *S. sp.* NRRL F-2890<br>*PermE* * <sub>pHM11a</sub> - *cftA* <sub>JV178</sub> | Undetected | 19.84 ± 4.86 μM | Undetected | 16.56 ± 0.01 μM |
| *S. negayawaensis* | Undetected | 19.53 μM | 6.46 μM | 7.20 μM |

To understand clifednamide biosynthesis towards engineering its production, JV178 was genome-sequenced (see Methods). Due to the structural similarity of ikarugamycin and clifednamides, we expected JV178 would encode an ikarugamycin-like PTM cluster. Using BLAST, we identified a 5-gene PTM locus that likely encodes the clifednamides. Homology analysis of these genes allowed us to propose a plausible clifednamide biosynthetic pathway (Figure S1).

Three of the genes, *ikaA*, *ikaB* and *ikaC,* recapitulate the ikarugamycin cluster^12^. As such, clifednamide biosynthesis likely begins with IkaA, an iterative polyketide synthase/non-ribosomal peptide synthase fusion protein. Biochemical studies of its ortholog in HSAF biosynthesis^17^ indicate the protein initiates PTM biosynthesis by ligating two polyketide chains, built from six malonyl-CoA precursor units each, to the non-proteinogenic amino acid L-ornithine (Figure S1)^18^. The resulting tetramate-polyene product is reductively cyclized by IkaB and IkaC to produce the 5-6-5 carbon ring system shared by ikarugamycin and clifednamides^19^. A *ftdA* homolog was also found in the cluster. A PTM hydroxylase common to a number of PTM pathways^7,20,21^, *ftdA* is likely responsible for the C25 hydroxyl group of clifednamide B. The remaining open reading frame is encoded between *ikaA* and *ikaB* (Figure 1A). Predicted to encode a novel cytochrome P450, we reasoned its cognate enzyme (designated CftA, for <u>c</u>lifednamide tailoring <u>A</u>) may install the C29 ketone of the clifednamides. No additional PTM genes were detected in the JV178 genome, further suggesting the *cftA*-containing cluster encodes the clifednamides.

JV178 was found to be a poor host for genetic analysis (unpublished), preventing experimental verification of the above model. To further study clifednamide biosynthesis, a genome mining approach was used to identify additional producers. Several actinomycetes having publicly available genome sequences were found to harbor PTM clusters syntenic with the JV178 locus (Figure S2). Four such strains were obtained from the USDA NRRL strain collection. Each strain was grown on a panel of solid media and extracted with ethyl acetate for LC-MS/MS analysis. Clifednamide production was determined by comparison with extracts containing clifednamides A and B from JV178. All four strains appear to produce both compounds based on product retention times, UV absorbance spectra, and mass fragmentation patterns (Figure 2). Notably, *S. negayawaensis* produced approximately 26 μM of clifednamides A and B combined, about 10-fold greater than JV178 (Figure 2E, Table 1). The other three strains, *S. purpeofuscus*, *S.* sp. F-6131, and *S. torulosus* produced considerably lower amounts (Figure 2B-D). Interestingly, a local PTM-producing soil isolate (*Streptomyces* sp. KL33) also produced clifednamide A titers comparable to *S. negayawaensis*, but no clifednamide B was detected (Figure 2F).

**Figure 2.**
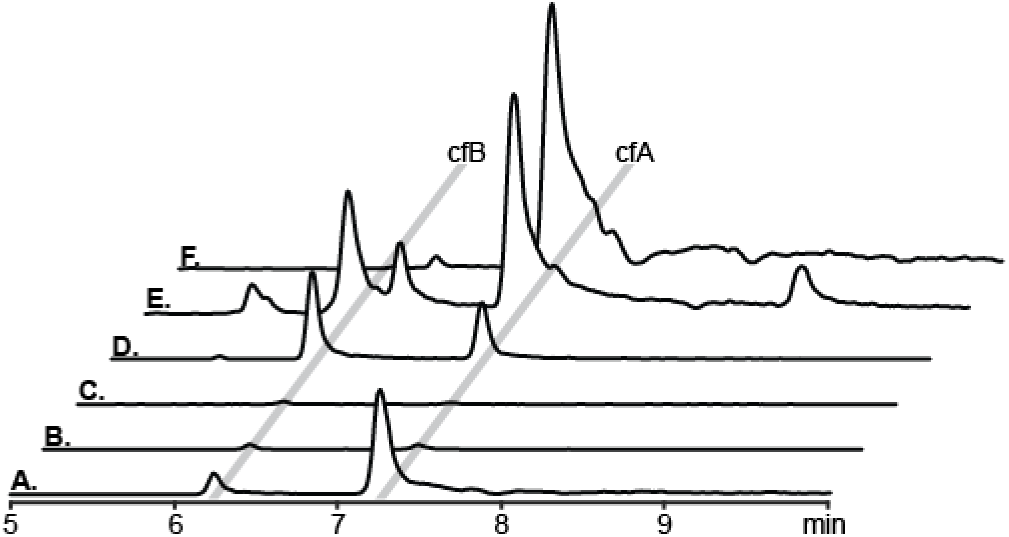
Clifednamide production by genome-mined strains. LC-MS/MS chromatograms of extracts from (A) JV178, (B) *S. purpeofuscus*, (C) *S.* sp. F-6131, (D) *S. torulosus*, (E) *S. negayawaensis*, and (F) KL33. The retention times for clifednamides A and B (cfA and cfB) are indicated by grey bars.

Of the newly obtained clifednamide producers, *Streptomyces*. *sp.* NRRL F-6131 proved to be the most amenable to intergeneric conjugation. This strain was thus used to interrogate clifednamide biosynthesis. A *rpsL*(K43N) mutant was isolated for streptomycin counterselection^22^. As noted for other *Streptomyces* metabolites, this lesion also increased PTM production^23^ (Figure 3A&B). Markerless *cftA* deletion resulted in the loss of clifednamide production. However, the strain produced increased amounts of a previously minor compound (*m/z* 479) (Figure 3C). This peak was confirmed to be ikarugamycin by comparison with an authentic standard, and its apparent accumulation is consistent with it being a clifednamide biosynthetic precursor. An additional peak of interest (*m/z* 495) was detected in the Δ*cftA* mutant. This was tentatively identified as butremycin, a known C25-hydroxyl derivative of ikarugamycin^24^. The experimental mass matches the compound and its UV profile is consistent with other PTMs. MS/MS analysis revealed a daughter ion with a *m/z* of 154, consistent with the C25-hydroxylated PTMs such as clifednamide B. Because butremycin is structurally equivalent to clifednamide B lacking the C29 ketone, it is an expected biosynthetic precursor.

**Figure 3.**
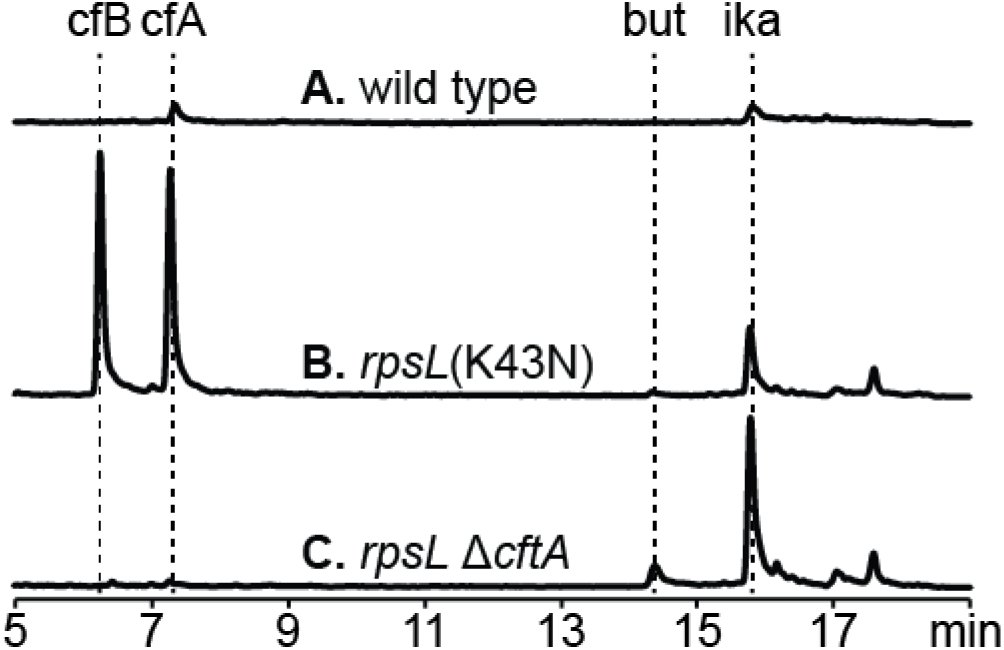
Deletion of *cftA* in *S.* sp. F-6131. LC-MS/MS chromatograms of extracts from (A) wild-type *S.* sp. NRRL F-6131, (B) the *rpsL* (K43N) mutant, and (C) the *rpsL* Δ*cftA* double mutant. The peaks for ikarugamycin (ika), butremycin (but), clifednamide A (cfA), and clifednamide B (cfB) are indicated with dotted lines.

Our biosynthetic analyses suggested an ikarugamycin-producing microbe could be engineered to produce clifednamide via *cftA* expression. Using genome mining, *Streptomyces* sp. NRRL F-2890 was identified as a candidate host. Characterization of the strain revealed it has robust flux, producing up to 28 μM (13.6 mg/L) of ikarugamycin (Figure 4A). A genetic system was established in the strain to systematically evaluate the biosynthetic effects of a panel of CftA isozymes. To do this, four *cftA* orthologs were cloned under two versions of the strong constitutive P*ermE*\* promoter. While the original *ermE* promoter in *Saccharopolyspora erythraea* begins transcription at the start codon^25^, a short 5’-UTR containing a ribosome binding site is often added to P*ermE*\* expression plasmids. The two plasmids used in this study have P*ermE*\* promoters with variant 5’-UTR sequences that differentially express a *xylE* reporter (Figure S3)^26^.

**Figure 4.**
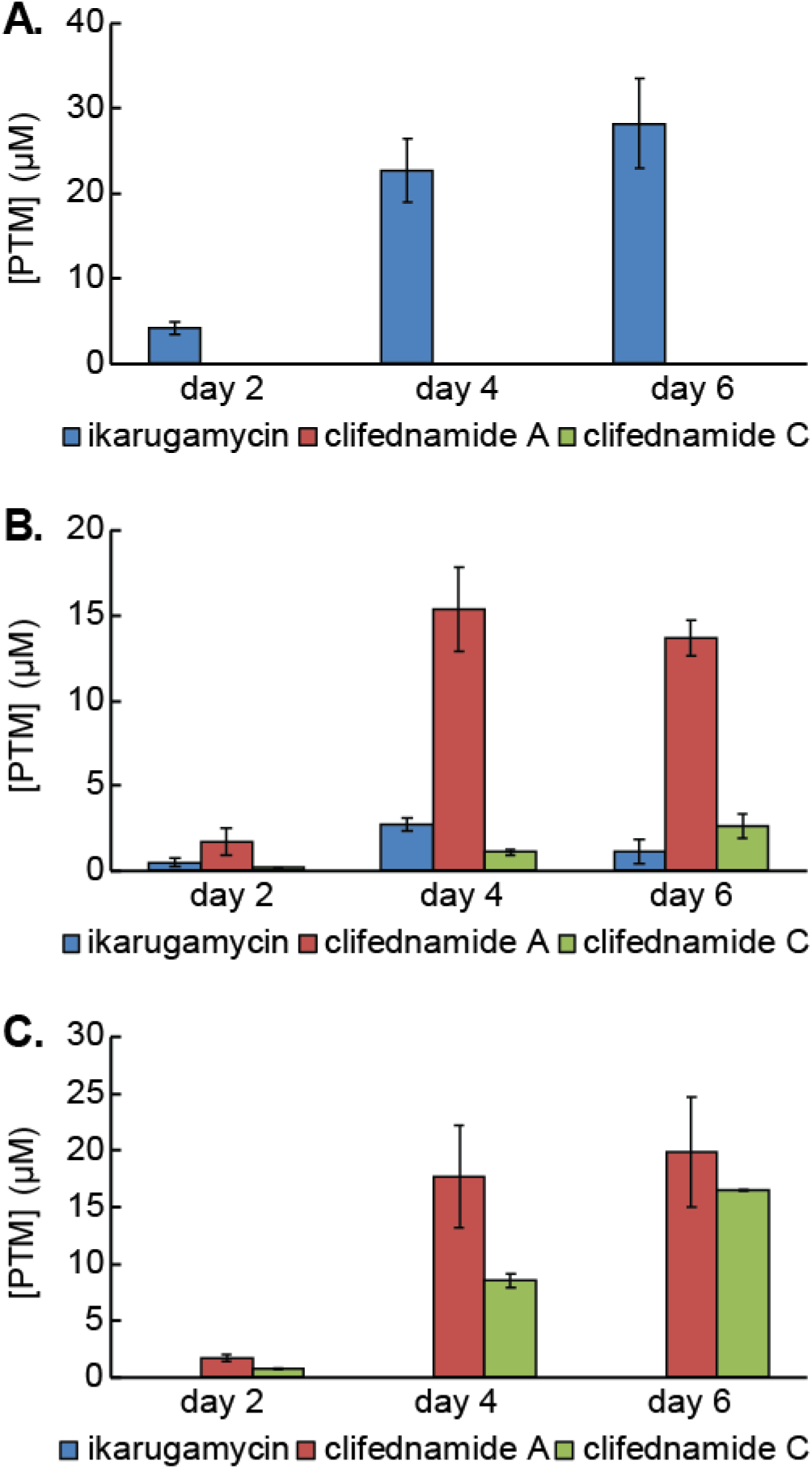
Production of PTMs in *Streptomyces* sp. F-2890 expressing the JV178 CftA. (A) Wild-type F-2890, (B) F-2890 *attB*ΦC31::P*ermE*\*_pDA1652_-*cftA*_JV178_, and (C) F-2890 *attB*ΦC31::P*ermE*\*_pHM11a_-*cftA*_JV178_. Strains were grown in triplicate and extracted at days 2, 4, and 6.

The resulting panel of P*ermE*\*-*cftA* constructs were chromosomally integrated in F-2890 at the ΦC31 *attB* site^27^ following intergeneric conjugation. As expected, the resulting strains all produced clifednamide A (Figures 4, S4). Interestingly, CftA homologs from different strains produced varying amounts of clifednamide A, with the JV178 homolog producing the most (20 μM, Table 1). No clifednamide B or butremycin was observed in any of these strains due to *ftdA* being absent in the host. In general, constructs driven by the stronger pHM11a^28^-derived promoter converted more ikarugamycin precursor to clifednamide A than the pDA1652-derived promoter^29^.

Additionally, all four pHM11a constructs and two pDA1652 constructs resulted in a new peak having a characteristic PTM UV profile and *m/z* 509 (Figures 4, S4). MS/MS fragmentation produced a daughter ion with *m/z* of 139, a diagnostic daughter ion of PTM molecules lacking C25 hydroxylation such as clifednamide A, ikarugamycin, and capsimycin^30^. Furthermore, metabolic labeling with ^2^H_7_-L-ornithine results in a correspondingly heavier mass (*m/z* 516) and daughter ion (*m/z* 146). These results parallel our results from clifednamide A (Figure S5). Based on these results, we designated this peak clifednamide C. Its mass difference (*m/z* +30) from ikarugamycin (*m/z* 479) and shorter C18 retention time suggest it has been oxidized twice. The first oxidation presumably corresponds to the C29 keto oxidation by CftA to produce clifednamide A (*m/z* 493). The subsequent oxidation of clifednamide A to clifednamide C is consistent with a hydroxylation (*m/z* +16), perhaps resulting from above-optimal CftA activity. A clifednamide C peak was also detected in *S. negayawaensis*, confirming its production in un-engineered clifednamide producers.

The tandem oxidation of clifednamide A by CftA is analogous to the activity of the recently characterized IkaD^30^. A cytochrome P450 associated with some ikarugamycin biosynthetic clusters, IkaD primarily installs an epoxide across carbons 7 and 8. However, it can also hydroxylate C29, the same position targeted for keto insertion by CftA (Figure S1). We noted that *Streptomyces* sp. NRRL F-2890, the ikarugamycin-producing host strain, also contains an IkaD homolog. While F-2890 produces a small peak that is consistent with the mass of ikarugamycin epoxide, this peak remains minor in CftA expression strains (unpublished data). Therefore, CftA apparently lacks the epoxidase activity of IkaD.

Presumed CftA orthologs mined from public sequence repositories all share at least 80% amino acid sequence identity, but have no more than 57% residue identity with the IkaD orthologs. Thus, under the cytochrome P450 naming convention^31^, CftA and IkaD are members of distinct subfamilies within the CYP107 clade. Aside from IkaD and CftA, additional PTM-locus P450 enzymes also fall within the CYP107 group (Figure S6, S7). Beyond CYP107 family P450's, our data also indicates the existence of three other distinct CYP families (<40% identity with CYP107) associated with genome mined PTM-loci. With the largest group being comprised of FtdF homologs (as found in the frontalamide cluster), no role has been established yet for these enzymes. Together, our analyses suggest further investigations into PTM-associated CYPs will continue to reveal enzymes with varied PTM scaffold specificity, oxidative activities and regioselectivity.

In conclusion, we used a combination of comparative genomics, genetics, and genome mining to reveal the novel cytochrome P450 CftA is responsible for clifednamide production. We engaged in a synthetic biological approach to engineer clifednamide production by utilizing *cftA* orthologs sourced from newly identified clifednamide producing strains. In addition to increased clifednamide production over *Streptomyces* sp. JV178, we noted substantial differences from *cftA* homologs. As such, this work demonstrates the value of leveraging genome-mined panels of isozymes in a plug-and-play fashion to rapidly identify efficient biocatalysts. Further, we show CftA belongs to an expanding family of PTM cytochrome P450 enzymes with the apparent capability to carry out multiple oxidation events. These enzymes may be useful to engineer diversity-oriented panels of PTMs for future bioactivity and structure-function analyses.

## METHODS

### Strains, Plasmids, Primers, Enzymes, Chemicals and General Methods

Strains, plasmids, and primers are described in Tables S1-3. Several strains were obtained from the Agricultural Research Service Culture Collection (NRRL). All primers were purchased from Integrated DNA Technologies. All restriction enzymes and Taq polymerase were purchased from New England BioLabs. T4 ligase was purchased from New England BioLabs and ThermoFisher. KOD Hot Start DNA Polymerase (EMD Millipore) in FailSafe PCR 2X PreMix (Epicentre) was used to amplify DNA sequences for cloning from *Streptomyces* genomic DNA. Taq polymerase was used for colony PCR. Ikarugamycin (>99%) was purchased from Santa Cruz Biotechnology. *L*-Ornithine-2,3,3,4,4,5,5-d_7_ HCl (>98% atom D) was purchased from CDN Isotopes. All other chemicals were obtained from Sigma Aldrich or Fisher Scientific. *Streptomyces* genomic DNA was prepared for PCR by grinding a colony in 100 μL DMSO as described by Van Dessel *et al*^32^. Standard protocols for manipulating *E. coli* were based on those of Sambrook *et al*^33^. Streptomycetes were routinely propagated on ISP2 agar^34^ and Trypticase Soy Broth (Difco) at 28°C. Glass beads were added to liquid cultures to disrupt mycelial clumps.

#### Genome sequencing of JV178

Genomic DNA was extracted from TSB-grown mycelia as previously described^35^. Illumina 250-bp paired-end sequencing libraries were prepared using the Nextera sample prep kit (Illumina Inc., San Diego, CA, USA) and were sequenced on an Illumina MiSeq platform using V2 chemistry (Illumina, Inc., San Diego, CA, USA) by the Washington University in St Louis Genome Technology Access Center. Sequencing reads were trimmed and *de novo* assembled using the CLC Genomics Workbench (CLC Bio-Qiagen, Aarhus, Denmark). An annotated sequence for the *Streptomyces* sp. JV178 clifednamide cluster was deposited to GenBank (accession no. MF89327).

#### Streptomycete conjugations

*S.* sp. F-6131 spores were collected from ISP4 agar^34^, while *S.* sp. F-2890 spores were collected from ISP2 agar. Spores were harvested using TX Buffer^36^. Conjugations were performed using JV156 as the general *E. coli* donor as previously described^7^. Exconjugants were selected with 50 μg/mL colistin and 25 or 50 μg/mL apramycin. Successful conjugations were verified by colony PCR.

#### *cftA* markerless gene disruption

The *cftA* coding sequence of *S.* sp. F-6131 was replaced with a truncated gene containing the first nine codons and the last ten codons of the wild-type coding sequence with homologous recombination as previously described^37^. Streptomycin-resistant (Str^R^) mutants of *S.* sp. F-6131 were isolated on ISP2 + Str^25^ agar. The *rpsL* genes were amplified and sequenced. JV739 bearing the *rpsL* (K43N) mutation was chosen for subsequent experiments as the mutation did not disrupt clifednamide production. The 990 bp upstream flanking region of *cftA* was amplified using primers YQ273 and YQ274 (introduced a *Xba*I site and homology to pUC19). The 1079 bp downstream flanking region of *cftA* was amplified using primers YQ275 (introduced homology to the upstream flanking region) and YQ276 (introduced a *Hind*III site and homology to pUC19). The 2668 bp fragment of pUC19 was amplified with primers YQ268 and YQ269. PCR amplicons were assembled using the NEBuilder HiFi Assembly kit (New England BioLabs). The resulting pUC19-Δ*cftA* was digested with *Xba*I and *Hind*III and the 1956 bp fragment was ligated into pJVD52.1 digested with *Xba*I/*Hind*III. The resulting pJVD52.1-Δ*cftA* was introduced into JV739 by intergeneric conjugation, and apramycin-resistant (Apr^R^) exconjugants were selected. Exconjugants were grown in TSB non-selectively at 37°C and double-recombinants were selected for on ISP2 + Str^100^. The Δ*cftA* mutants were confirmed by PCR.

#### *cftA* heterologous expression

The *cftA* homologs were amplified using primers ED9-16 (introduced *Nde*I and *Xba*I sites). The PCR products were digested with *Nde*I and *Xba*I and ligated into pJMD2 or pJMD3 to generate plasmids pED1-8. After confirming the inserts by Sanger sequencing (Genewiz), the constructs were introduced into *S.* sp. F-2890 by intergeneric conjugation.

#### PTM detection by HPLC-MS/MS

Strains were cultivated in 15 mL of TSB liquid medium in 125 mL Erlenmeyer flasks shaken in 1-inch orbitals at 250rpm at 28°C. 6 mm glass beads were added to disrupt mycelial clumps. After 2 days of growth, 200 μL of cultures were plated on HT^38^, ISP4, ATCC172, or JBFM1 (adapted from Medium 2^39^: 2% D-fructose; 5% D(+)Mannose; 0.167% Na-L-aspartate; 0.06% L-arginine HCl; 0.05% L-histidine HCl; 0.2% K_2_HPO_4_; 0.2% KH_2_PO_4_; 0.5% NaCl; 0.006% ZnSO_4_-7H_2_O; 0.0256% MgSO_4_-7H_2_O; 0.051% MgCl_2_-6H_2_O; 0.001% CoCl_2_-6H_2_O; 0.036% NaSO_4_; 2.13% MES free acid; 1.5% agar; 2% R2 Trace elements; 1.5% Agar; pH 6.0) and incubated at 28°C. After 6 days, the agar was diced and immersed in ethyl acetate overnight. The ethyl acetate was evaporated at low pressure and the extract was suspended in 400 μL of HPLC-grade methanol and syringe filtered.

Analysis was performed using a Phenomenex Luna C18 column (75 x 3 mm, 3 μm pore size) installed on an Agilent 1260 Infinity HPLC connected to an Agilent 6420 Triple-Quad mass spectrometer using the following method: *T* = 0, 5% B; *T* = 3, 40% B; *T* = 13, 60% B; *T* = 17, 100% B, *T* = 20, 100% B; A: water + 0.1% formic acid, B: acetonitrile + 0.1% formic acid; 0.9 mL/min. 10 μL of the methanol-dissolved extract was injected per run. The precursor ion scan mode was used to identify molecules that fragmented (collision energy, 30 V) into daughter ions with m/z of 139.2 or 154.2. The resulting data was analyzed offline with Agilent MassHunter Qualitative Analysis software. PTMs were quantified using integrated peak areas of absorbance at 320nm detected with an in-line diode array detector (DAD). A standard curve was generated using an authentic ikarugamycin standard.

## ASSOCIATED CONTENT

Supplementary methods, tables, figures, and references

## AUTHOR INFORMATION

### Author contributions

Y.Q., E.D., and J.A.V.B. designed the experiments and wrote the manuscript. Y.Q. and E.D. performed experiments.

### Notes

The authors declare no competing interest

## ACKNOWLEDGEMENTS

We thank Dr. Arpita Bose for helpful comments and discussions and John M. D’Alessandro for plasmids pJMD1, pJMD2, and pJMD3. We acknowledge former WUSTL BIOL3493 students Kevin Lou for isolation of strain *Streptomyces sp.* KL33 and Naveen Jain for designing *attB*^ΦC31^ integration check primers. We also thank Dr. Kim Medley of Tyson Valley Research Center for access to soil samples used in the isolation of KL33. This work was supported by Washington University in St Louis New Faculty Start Up funds to Joshua Blodgett.

---

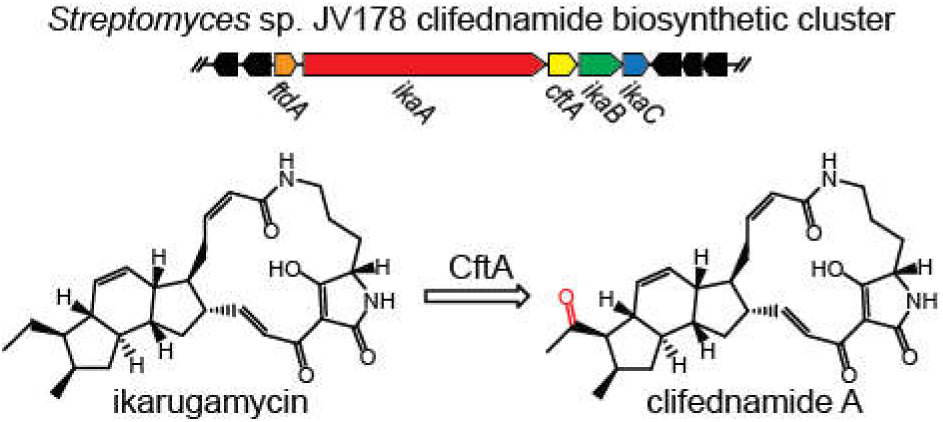
Graphical abstract

